# Channel crosstalk correction in suspension and imaging mass cytometry

**DOI:** 10.1101/185744

**Authors:** Stéphane Chevrier, Helena Crowell, Vito Zanotelli, Stefanie Engler, Mark D. Robinson, Bernd Bodenmiller

## Abstract

Mass cytometry enables simultaneous analysis of over 40 proteins and their modifications in single cells through use of metal-tagged antibodies. Compared to fluorescent dyes, the use of pure metal isotopes strongly reduces spectral overlap among measurement channels. Crosstalk still exists, however, caused by isotopic impurity, oxide formation, and mass cytometer properties. Spillover effects can be minimized, but not avoided, by following a set of constraining rules when designing an antibody panel. Generation of such low crosstalk panels requires considerable expert knowledge, knowledge of the abundance of each marker and substantial experimental effort. Here we describe a novel bead-based compensation workflow that includes R-based software and a web tool, which enables correction for interference between channels. We demonstrate utility in suspension mass cytometry and show how this approach can be applied to imaging mass cytometry. Our approach greatly simplifies the development of new antibody panels, increases flexibility for antibody-metal pairing, improves overall data quality, thereby reducing the risk of reporting cell phenotype and function artifacts, and greatly facilitates analysis of complex samples for which antigen abundances are unknown.

## INTRODUCTION

High-dimensional, single-cell flow cytometry has been broadly adopted by researchers and clinicians to analyze complex biological samples^1–6^. Fluorescence activated cell sorting (FACS) has dominated this field for decades, and, with the constant improvement of probes and laser systems, 18-color FACS experiments are now routine^7^, and 30-color cytometers have recently become commercially available^7^. Due to the overlapping excitation and emission spectra of the fluorescent dyes, signals are measured not only in the primary channel, but also in neighboring channels. This spillover is correlated with original signal in an approximately linear manner and can be corrected via a process called compensation^8^. Single-stained controls for each dye are analyzed to determine the percentage of interfering signal in all the channels. These values are reported into a spillover matrix that is then used to correct for the spillover by solving a system of linear equations. As the number of parameters measured increases, however, it becomes challenging to detect proteins of low abundance due to the increased complexity of overlapping spectra of fluorochromes that interfere with detection in the affected channels^7^.

The recent development of mass cytometry increased the number of parameters that can be analyzed in an experiment to over 40^9^. In mass cytometry, heavy metal isotopes are used as reporters to label antibodies^10–12^. Currently, mass cytometers enable simultaneous detection of 135 channels ranging from 75 to 209 atomic mass units. To date, more than 55 different isotopes have been used for surface and intracellular protein epitope detection, DNA and transcript detection, cell-cycle analysis, dead-cell exclusion, and cell barcoding^2,13–16^. This technology has recently been exploited for imaging by coupling a laser ablation system to a mass cytometer^17–19^. Imaging mass cytometry (IMC) enables the analysis of tissue sections stained with metal-tagged antibodies to generate highly multiplexed images at subcellular resolution^17,18^.

Due to the high mass resolution of the time-of-flight mass spectrometer and the fact that the ions are singly charged and thus differ by at least a full atomic mass unit, the interference between channels in mass cytometry is strongly reduced in comparison to the overlap observed in fluorescent flow cytometry^20^. Still, three sources of signal interference are observed in mass cytometry^11^. The first is imprecision in ion detection (at mass M) caused by small differences in ion positions and kinetic energy at the beginning of the mass analysis process leading to signal leakage into adjacent mass channels (M±1). This source of spillover is termed abundance sensitivity and is a fixed value for each instrument under a specific configuration. In current mass cytometers, the (M±1)/M ratio is low and optimized to not exceed 0.3% for ^159^Tb^21^. The second source of signal interference is oxide formation, which generates a signal in the M+16 channel. The frequency of oxide formation after ionization depends on the strength of the oxygen-ion bond and the temperature of the plasma. The tuning procedure for mass cytometers ensures that the oxide ratio remains under 3% for lanthanum (^139^La), an easily oxidized isotope. The third source of spillover is due to the impurity of the isotopes used in mass cytometry. For relatively abundant isotopes, the isotopic purity is usually greater than 99%, whereas for less abundant isotopes, impurities in a single alternative channel can reach 4%^11,20^.

Although the amount of spillover observed in mass cytometry is generally small, spillover can considerably complicate interpretation of data and potentially lead to false conclusions. For example, signal crosstalk can result in incorrect identification of cells as expressing an intermediate level of a marker^22^. In experiments conducted to date, the effects of spillover have been minimized by selecting only highly pure isotopes and by carefully designing antibody panels to optimize the signal to background ratio in each channel^22^. Generating a low crosstalk antibody panel is complex and time consuming, however. It requires that the approximate antigen abundance is known for each marker used in the panel, which is not possible in many types of experiments, particularly when a large variability of expression levels of a particular marker is expected or when the expression levels can unpredictably increase through an applied perturbation or a disease state. Thus, especially clinically important samples, such as complex tissue-derived samples, can suffer from spillover artifacts. Further, with a purely experimental approach to avoid spillover, antibody-isotope conjugates are not easily transferable between panels applied to different sample types. As spillover is proportional to the originating signal, it can be reduced by decreasing antibody concentrations; this also reduces the signal-to-noise ratio and is thus only applicable to channels with clear positive and negative populations. In practice, the above-mentioned strategies are not sufficient to completely prevent crosstalk between channels as shown in a recent study in which data from spillover-affected channels had to be excluded to avoid potentially misleading conclusions^23^. Spillover-related issues have not yet been reported in IMC, but since the source and the measurement of metal signal in suspension mass cytometry and IMC are identical, both systems are expected to be affected in a similar manner.

Here we present a comprehensive workflow to estimate and systematically correct for signal spillover across all the channels used in a given mass cytometry experiment. Polystyrene capture beads were single-stained with each antibody used in the experiment. Beads were then pooled and analyzed simultaneously in the mass cytometer. The CATALYST R/Bioconductor package and an interactive Shiny-based web application were developed to deconvolute the different bead populations, estimate spillover signal in all channels, and compensate the data. We demonstrate the utility of the approach in correction of signal interference in suspension mass cytometry and IMC experiments. Our approach will greatly facilitate the development of antibody panels, increase the flexibility of antibody-metal pairing, increase the number of usable isotopes, and enable generation of high-quality data void of spillover artifacts on poorly characterized samples.

## RESULTS

### Mass cytometry spillover is linear and can be corrected using compensation

Fluorescent flow cytometry is affected by signal interference between channels. Since spillover signal is a defined fraction of the source signal, it can be corrected mathematically^8,24^. In mass cytometry, the interference between channels is reduced but is still present due to instrument properties (abundance sensitivity), isotopic impurities, and oxidation (Figure 1A). To determine whether channel crosstalk observed in mass cytometry can be corrected in a manner similar to the one used for flow cytometry, we first determined whether the crosstalk in mass cytometry experiments is linear. We stained peripheral blood mononuclear cells (PBMCs) with anti-CD44 conjugated to ^143^Nd using antibody concentrations ranging from 0.01 μg/mL to 1 μg/mL (Figure 1B). As expected, signal was observed in other mass channels including −1 (^142^Nd), +1 (^144^Nd), +2 (^145^Nd), +3 (^146^Nd), and +16 (due to the oxidation product ^143^Nd^16^O measured in ^159^Tb). The signal in the source and in the spillover channels could be fitted by a linear model with a Pearson correlation coefficient (R) greater than 0.99 in all cases (Figure 1C).

**Figure 1.**
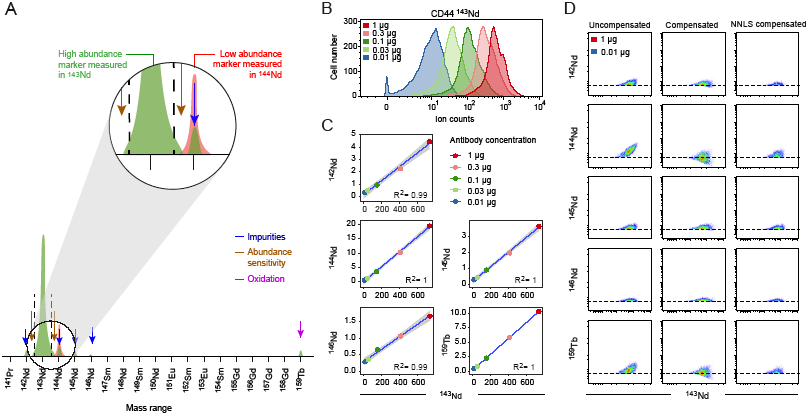
Spillover is a linear function of the original signal and can be compensated. **A.** Schematic view of sources of signal interference affecting mass cytometry. **B.** Histogram showing signal intensity upon staining of PBMCs with the indicated concentrations of anti-CD44 antibody. **C**. The median intensities of signals obtained in the main channel (^143^Nd) and spill-affected channels for PBMCs stained with anti-CD44 antibody are displayed as scatter plots. For each relationship, the Spearman coefficient of correlation is indicated. **D**. Scatter plots showing the signals of the anti-CD44 antibody in the main channel and in the spillover-affected channels before compensation (left column), after compensation with the conventional fluorescent flow cytometry approach (middle column), and after compensation with the NNLS method (right column).

In fluorescent flow cytometry, the slope of the linear regression is used to generate a spillover matrix. Flow cytometry compensation approaches apply this spillover matrix by solving a linear system of equations to infer the original signal values. Applying the traditional flow cytometry compensation on these single-stained cells efficiently removed the spillover (Figure 1D, middle panels). However, this strategy substantially modified the structure of the data by introducing artificial negative values, which specifically influenced channels strongly affected by spillover. Negative ion counts are not present in uncompensated mass cytometry data, and more importantly, data with negative values require different treatment than strictly non-negative abundance data. A recent study aimed at unmixing signals in multispectral fluorescent flow cytometry made similar observations and suggested use of approaches that specifically incorporate a non-negativity constraint for the compensation process, such as the ‘Non Negative Least Squares’ (NNLS) approach^25^. This method calculates the optimal non-negative solution for the compensation problem using the least-squares criterion. Applied to our data, the NNLS approach removed the spillover without changing the data structure (Figure 1D, right panels). Taken together, spillover in mass cytometry is linear and can be corrected while preserving the data structure using the NNLS approach.

### Systematic correction of spillover in mass cytometry

Inspired by methods used in flow cytometry, in which controls stained with single antibodies are used to estimate signal crosstalk, we developed an approach to systematically correct for signal interference in mass cytometry experiments. A 36-antibody panel was designed to detect the main immune cell populations in PBMCs (Figure 2A). This panel was not optimized to avoid spillover effects. We also labeled identical antibodies for detection in different mass channels to facilitate the identification of spillover artifacts and to validate our method. In parallel to sample staining with the panel, 36 control samples stained with individual antibodies were generated by staining polystyrene antibody-capture beads in a 96-well plate (Figure 2B). After staining, beads were pooled and run as a single sample in the mass cytometer.

**Figure 2.**
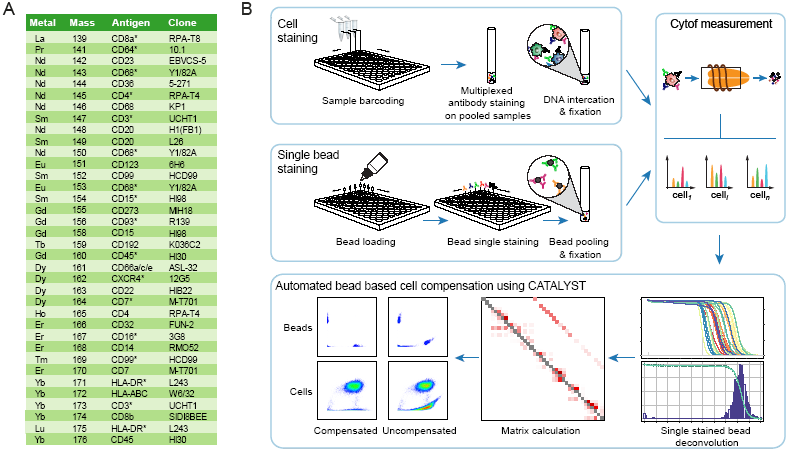
Antibody panel and experimental and analytical workflow to correct spillover using single stained beads. **A.** List of the 36 antibodies used in the panel in this study, including the information regarding the metal, the mass, the antigen, and the clone. Asterisks indicate antibody grouping for the experiment shown in Figure 3C. **B.** Depiction of the workflow used to correct for spillover. Staining of control antibody-capture beads and samples are performed in parallel. Single stained beads are pooled, and data are acquired on the beads and the samples by mass cytometry. The CATALYST R package enables identification of the single-positive bead populations, calculates the compensation matrix, and applies the matrix to correction of sample data for spillover.

To apply our approach for semi-automatic spillover correction in mass cytometry, we created an R/Bioconductor package, CATALYST, and a web app (Figure S1A). In the first step, the FCS file containing data on the bead sample is deconvoluted to identify the individual single-antibody-positive bead populations using a new implementation of the debarcoding algorithm from Zunder and colleagues^114^ that includes automated estimation of sample-specific cutoffs (Figure S1B). In a second step, the spillover matrix is calculated based on the spillover observed for single-stained populations. Due to the mass cytometry data structure, we observed that spillover estimation was more accurate when the spillover was assessed at the single bead level rather than at the bead population level (Figure S2A - C; see Materials and Methods for details). By default, the method only takes into account interference between channels expected to interact based on abundance sensitivity, metal impurity, and oxidation (Figure S2D, E), but also allows to check for unexpected spillover. In a final step, the compensation matrix from the solved linear system (NNLS or classical) is applied to the bead and cell samples to remove interfering signal. This workflow provides a fully integrated and easy to use experimental and computational solution for compensation of mass cytometry spillover.

### Cellular metal load influences signal spillover

The spillover matrix generated by our bead approach revealed that the total amount of spillover originating from a single channel ranged from 0% for ^165^Ho to over 8% for ^148^Nd (Figure 3A). This spillover matrix also revealed important differences regarding the amount of spillover received by the different channels. Consistent with previous reports^20^, we found that oxidation was variable among the different metals, ranging from 0% to 2%, but that the spillover due to mixed effects of impurity and abundance sensitivity is most problematic and may exceed 4%. Signal interference due to abundance sensitivity alone was virtually absent on the machine used. To assess the stability of the spillover matrix over time and instruments, we measured single-stained beads on different mass cytometers over months (Figure S3). Although the spillover matrix could be stable over months, our results show that for optimal compensation the spillover matrix should be acquired simultaneously with the sample of interest.

**Figure 3.**
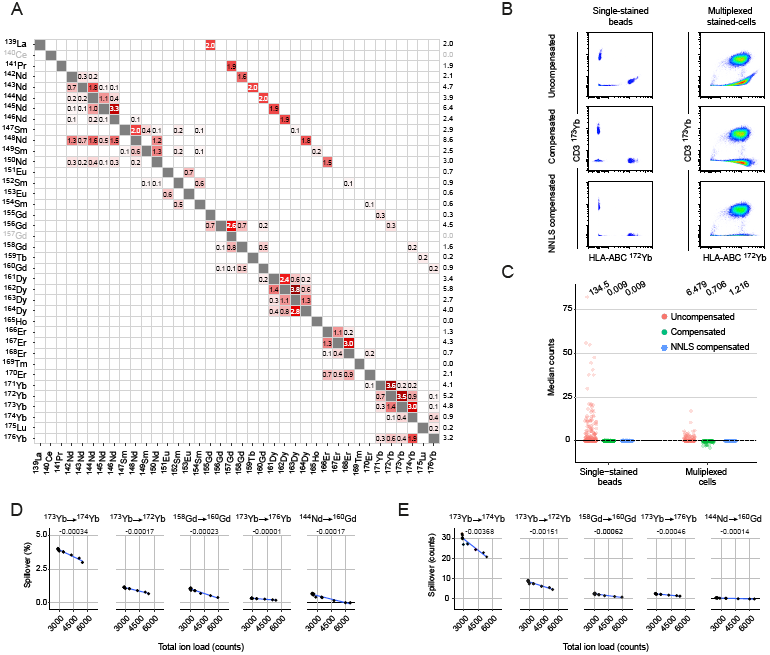
Saturation effects cause overcompensation in multiplexed samples. **A.** Spillover matrix calculated based on single-stained beads. Values on the diagonals are one. By default, spillover is calculated only in potentially affected channels, which include M±1, those corresponding to known isotopes, and M+16 (Figure S2D). Numbers in the cells indicate percentages of spillover by channels in rows into channels in columns. Numbers in the last column show the total amount of signal received in the corresponding channels. **B.** Scatter plots showing signal due to anti-HLA-ABC labeled with ^172^Yb and anti-CD3 labeled with ^173^Yb from pooled single-stained beads and multiplexed-stained PBMCs before and after compensation. The last row shows NNLS compensated data. **C.** Dot plots showing the median counts in each channel potentially affected by spillover for uncompensated data, compensated data, and NNLS compensated data obtained upon analyses of single-stained beads and multiplexed-stained PBMCs. For multiplexed-staining, cells were stained with two panels where half of the channels were left empty (see Figure 2A), to enable spillover assessment in absence of staining. For each dataset, the average sum of squares is shown on top of the graph. **D.** Dot plots showing the spillover in percent for the indicated relationships assessed on cells stained with increasing amount of barcoding reagents. A linear model was fitted to each relationship (blue lines), and the slope is indicated above each plot. **E.** Dot plots showing the spillover in absolute counts for the indicated relationships assessed on cells stained with increasing amount of barcoding reagents as described in panel D.

As expected, the application of the spillover matrix to beads stained with individual antibodies revealed virtually perfect compensation using both traditional and NNLS approaches (Figure 3B and C, left panels). When this matrix was applied to the multiplexed-stained cell samples, the spillover was also efficiently compensated but traditional compensation systematically overcompensated (Figure 3B and C, right panels). One possible explanation for the difference observed in spillover among single stained beads and multiplexed-stained cells might be the difference in total ion load, with high ion loads leading to ion detector saturation effects. Indeed, we found that an increased amount of barcoding was associated with a progressive decrease of spillover, both in terms of percentage and absolute count (Figure 3D, E). For spillover below two counts the signal interference was completely abolished (Figure 3E). While this observation leads to a slight but systematic overestimation of the spillover present in multiplexed stained cells, the NNLS compensation is robust to this effect and preserves the count abundance nature of mass cytometry data.

### Compensation efficiently corrects for spillover-mediated artifacts in mass cytometry

To identify spillover-based artifacts in our mass cytometry dataset and to determine the ability of our approach to correct them, we analyzed PBMCs stained with the 36-antibody panel using the dimensionality reduction algorithm t-SNE^26,27^. This approach enabled us to identify the main immune cell populations based on individual marker expression (Figure S4A). The proteins CD3, CD8, and HLA-DR were each detected with antibodies paired to two different metal-labels. In uncompensated data, we observed strikingly different signal profiles that depended on the metal isotope used to label the antibody (Figure 4A, left panel). After compensation, spillover signals were removed and the signals observed for the same antibodies conjugated with different metal isotopes were virtually identical (Figure 4A, right panel and Figure S4B). This demonstrated that compensation efficiently removed artifactual signals in spillover-affected channels and therefore prevented data misinterpretation during t-SNE map visualization.

**Figure 4.**
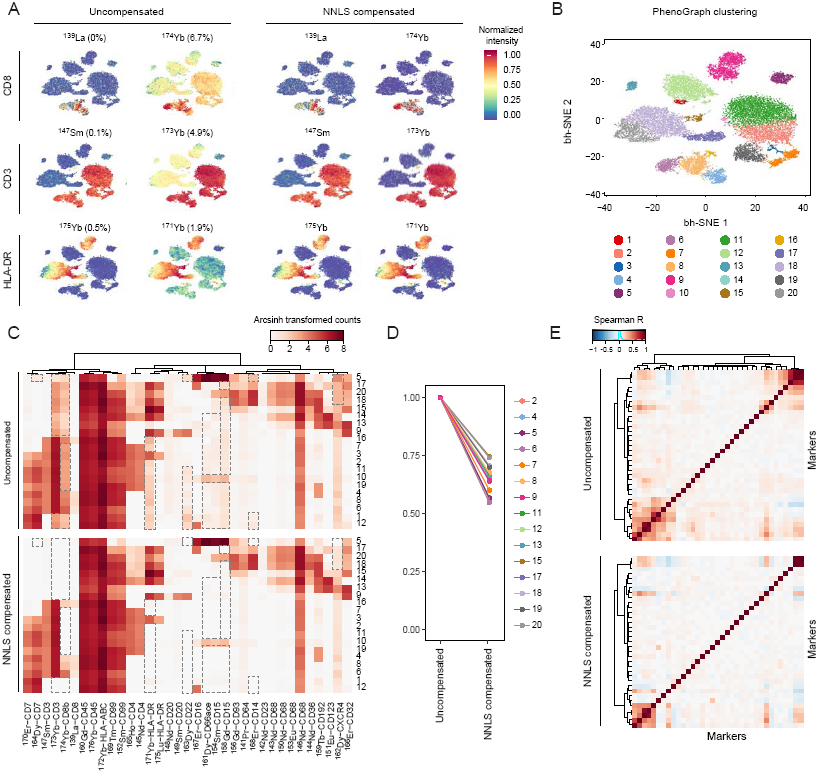
Correction of spillover artifacts in mass cytometry data using compensation. **A.** t-SNE map displaying data on a subset of 20,000 PBMCs analyzed with our 36-antibody panel and colored by marker expression for three pairs of antibodies labeled with two different metal isotopes before (left) and after (right) spillover correction based on NNLS compensation. The percentages of spillover affecting each channel in the uncompensated dataset are indicated. **B.** t-SNE map colored by PhenoGraph clusters identified on uncompensated data. **C.** Heat maps showing the expression of the indicated markers in the different clusters before compensation (upper panel) and after NNLS compensation (lower panel). Dashed boxes highlight regions in the plot that changed upon compensation. **D.** Plots showing the frequency of significant correlations (Spearman, p<0.005) between markers for each cluster containing more than 200 cells. Frequency was set to 1 for the uncompensated values. **E.** Correlation heatmap across all markers for cluster 12 before (upper panel) and after NNLS compensation (lower panel).

Identifying cell communities based on unsupervised clustering and investigating the marker expression profile of each population is an approach commonly used to analyze mass cytometry data. Applying the PhenoGraph^1^ clustering algorithm to our dataset led to the identification of 20 PBMC subsets (Figure 4B). Comparison of heat maps of signals in uncompensated versus compensated data highlighted how marker expression signatures of the different clusters can be misinterpreted without spillover correction (Figure 4C). Lack of compensation caused several clusters to be wrongly identified as having intermediate abundances of certain antigens even though the signal was actually due to channel crosstalk. In particular, an intermediate level of CD3-^173^Yb was observed on all the non T cell subsets (Figure 4C). Further, most T cell and natural killer cell subsets are wrongly identified as expressing intermediate levels of HLA-DR-^171^Yb. Artifacts caused by crosstalk were particularly strong in channels 158, 163, 168, 171, 173, and 174.

Characterization of newly identified clusters or signaling network inference often involves the systematic correlation analysis of markers at the single-cell level to identify co-regulated markers, and this approach can be strongly affected by channel interference. Analysis of marker correlations within each cluster before and after compensation systematically reduced spurious correlations (Figure 4D-E). A systematic analysis over all the clusters showed that in our dataset, between 25 and 45% of the significant correlations were actually due to spillover (Figure 4D). Collectively, this set of analyses showed that spillover can be responsible for various artifacts and that our compensation approach efficiently removed them.

### Spillover observed in IMC can be corrected using a similar compensation approach

In IMC, tissue stained with metal-tagged antibodies is ablated with a laser, and the tissue aerosol is analyzed in a mass cytometer^17^. Images generated with the IMC system provide subcellular resolution and are high dimensional; information has been collected from 50 different channels^17,18^. To determine how signal interference affects IMC measurements, metal isotopes were arrayed on a slide and measured by IMC. Using this approach, we demonstrated that a linear relationship exists between the original signal and the interfering signals (Figure 5A). This indicated that spillover in IMC could be corrected using the bead-based compensation approach applied to suspension mass cytometry. We used the CATALYST package to calculate a spillover matrix based on the pixel values of the individually spotted heavy metals (Figure S5A). Comparing individual spillover values obtained in suspension and IMC, we found that spillover due to abundance sensitivity and impurities were in the same range for all metals except for ^148^Nd and ^176^Yb, which came from different batches for the IMC experiment than those used for the suspension analysis (Figure S5B-D). Values observed for oxidation in the M+16 channel were systematically lower in IMC than in suspension mass cytometry. This was expected given that the tissue aerosol is transported in an argon and helium gas stream and no water is used for sample introduction, thus much less oxygen is present in the plasma of the mass cytometer to generate oxides (Figure S5E).

**Figure 5.**
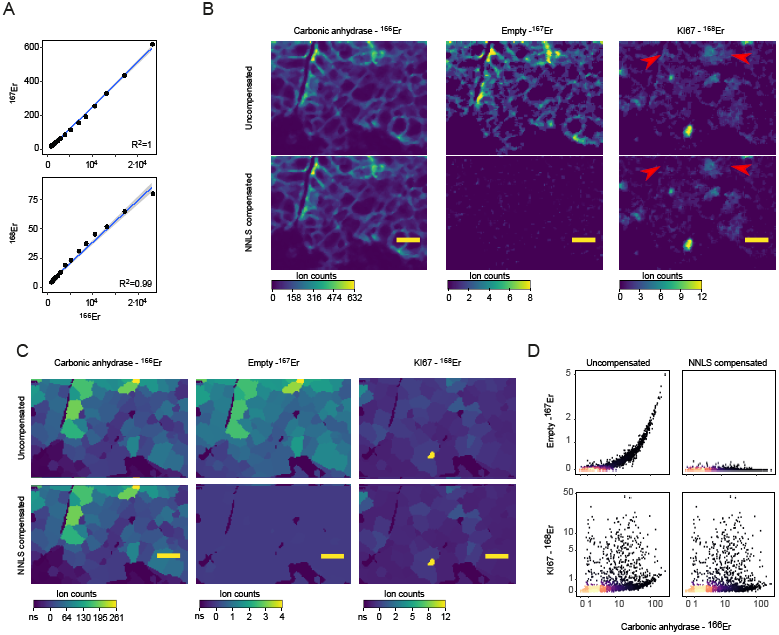
Spillover affects IMC data and can be corrected using our compensation strategy. **A.** Binning the signals of an imaged ^166^Er metal spot (to the 95^h^ percentile of the ^166^Er pixel values) into 20 bins with equal pixel numbers shows a linear relationship between ^166^Er and ^167^Er over several orders of magnitude (upper panel). The relationship between ^166^Er and ^168^Er appears linear but saturates at the higher counts (lower panel). **B.** Representative image of a breast cancer tissue sample imaged by IMC. Top row shows uncompensated images of ^166^Er (used to label antibody to carbonic anhydrase), ^167^Er (no antibody labeled with this metal), and ^168^Er (used to label anti-KI67). The bottom row shows corresponding NNLS compensated images. For visualization, a median filter was used to reduce noise. The yellow lines correspond to 20 um. Red arrows indicate part of the image where low signal was removed by compensation. **C.** Segmentation mask shown on representative images described in panel B. The mean pixel intensities of the signals observed in the indicated channels per cell are displayed. **D.** Scatterplots from single cell segmentation data from the IMC images before (left) and after (right) compensation with CATALYST. Arcsinh transformed ion counts (cofactor of 2) are shown.

Based on this spillover matrix, a breast cancer tissue section imaged by IMC was compensated at the pixel level using a custom written CellProfiler module^28^. This compensation approach efficiently removes the low signal due to spillover (Figure 5B). The carbonic anhydrase antibody, which was labeled with ^166^Er, showed a predominantly membranous signal. No antibodies were labeled with ^167^Er (the +1 channel) to enable assessment of the spillover. In the uncompensated data, there was a perfect, but lower intensity image of the ^166^Er channel in the ^167^Er channel. Thus, without compensation, the channel ^167^Er is not suitable for detection of a low level marker. ^168^Er was used to label an antibody against Ki67, a protein tightly regulated during cell cycle progression. Even though the spillover from ^166^Er into ^168^Er is only estimated to be 0.2% due to the low background in IMC, the carbonic anhydrase signal is still clearly visible in the KI67 channel. This could lead to the misinterpretation that Ki67 is localized on the membrane. Upon compensation, the shadow images of the ^166^Er channel in the ^167^Er and ^168^Er channels are gone (Figure 5B).

IMC images are often segmented to identify individual cells in the images enabling single cell data analysis^17,19^. Upon segmentation, the mean signal intensities for the channels of interest were calculated using a customized CellProfiler module and subsequently analyzed and compensated in R using the CATALYST package (Figure 5C). A scatter plot of this image clearly shows the spillover artifact and how our compensation approach largely removed it (Figure 5C). Together, this set of data indicates that spillover observed in IMC data can be and should be corrected using the compensation approach developed for correction of suspension mass cytometry data.

## DISCUSSION

Relative to fluorescence leakage in flow cytometry, channel interference is considerably reduced in mass cytometry, but it is not absent. Two reports have highlighted the challenges posed by spillover to mass cytometry data analysis and interpretation^22,23^. Although issues related to channel interference in IMC analyses have not yet been reported, our observations show that imaging and suspension mass cytometry are similarly affected by signal interference between channels. The main issue is the virtually absent background signal in mass cytometry, which enables the reliable detection of signals at low counts (~10 counts). Given the dynamic range over four orders of magnitude in mass cytometry, even a few percent signal spillover from high ion count channel into low ion count channel can easily result in difficulties in data interpretation. Moreover, high-dimensional mass cytometry data are commonly analyzed using unsupervised approaches, which present many advantages but involve a risk of misinterpretation of the data due to such artifacts. Currently, mass cytometry is transitioning from an emerging to a well-established technology, and this step requires the development of common standards and improved data quality assessment. Here, we present a combined experimental and computational approach that can be used both in suspension and IMC to ensure accurate correction of signal spillover.

In this study, we performed a comprehensive analysis of channel interference and showed that spillover is a linear function of the primary signal and therefore can be corrected using signal compensation similarly to flow cytometry. However, we found important differences between flow cytometry and mass cytometry data that prevented a direct transposition of the method used in flow cytometry. First, we found that assessing the spillover coefficient in single-stained beads using summary statistics at the population level tended to overcompensate the single-stained controls. Second, we observed that traditional compensation, by introducing negative values, changes the structure of the data. This artifact is not critical in fluorescent flow cytometry data as there are negative values in uncompensated data, but it has important consequences in mass cytometry data, in which negative events do not exist, but only events with zero or very low counts. The presence of physically impossible negative counts also changes the statistical properties of the data, as data can no longer be interpreted as abundances. Although not widely acknowledged, this problem has been already addressed in flow cytometry compensation methods used in multiparametric data analysis^25^. Third, we observed that the usage of a compensation matrix based on single-stained beads tended to overcompensate the spillover observed in multiplexed-stained cells, likely due to a detector saturation effect observed at higher ion counts.

The compensation method used in flow cytometry had therefore to be adapted to address these issues. To generate a more accurate compensation of the single-stained controls, we introduced a spillover estimation at the single-cell level, which was adapted to the mass cytometry data distribution and led to a perfect compensation of single-stained controls. To prevent appearance of negative data upon compensation, we applied the nonlinear least square method, previously used in multispectral fluorescent cytometry data compensation^25^. We found that this strategy preserved the structure of mass cytometry data. Finally, the overcompensation issue observed in multiplexed samples using the conventional compensation approach was virtually absent when using the NNLS approach, which strengthens the rationale for using this method.

Using a well-controlled system where the same antibodies were used in channels affected and not affected by spillover, we demonstrated that our approach efficiently removed spillover artifacts in t-SNE and PhenoGraph analysis and when investigating marker correlation within a given cluster. In straightforward experiments, most of the artifacts we describe here would have been detected by an expert user and removed by changing the antibody panel. In a complex experimental setup involving a large-scale analysis of poorly characterized tissues such artifacts are almost unavoidable, however, and may have gone unnoticed.

Compensating a ~40-parameter experiment can be seen as a daunting task. Therefore, the process was streamlined by using beads instead of cells for single-stained controls and by developing an efficient computational workflow that allowed for the simultaneous acquisition of data on all single-stained bead samples. All necessary data on the single-stained controls were acquired in a few minutes and were algorithmically identified as single-positive populations; these were subsequently used to calculate the matrix using the R package CATALYST, which includes a browser-based graphical user interface. For a 36-parameter imaging experiment, each individual metal can be acquired in less than a minute and upon acquisition, the samples can be processed by applying the CATALYST package on pixels instead of beads. The spillover matrix can be directly applied to the samples of interest or exported to be applied in commercial or open-source solutions such as FlowJo, Cytobank, or the R package flowCore for suspension experiments or a newly available plugin in CellProfiler for imaging experiments.

Mass cytometry is now broadly used in the scientific community. Our bead-based compensation workflow, including R-based software and a web tool, will make mass cytometry more reliable and easy to use. Minimizing spillover by careful design of the antibody panel is still advantageous, but our approach will offer a new level of flexibility, which is particularly needed for the analysis of complex and poorly characterized tissues such as tumor samples.

## DATA AVAILABILITY

Upon publication, all raw mass cytometry data (.fcs files) can be downloaded from http://www.bodenmillerlab.org and from the Cytobank repository. The R package developed in this study is available from Bioconductor (http://bioconductor.org/packages/CATALYST). The data processing pipeline can be run at the command line. Alternatively, an interactive shiny-based app can be run as either a local version (requires installation of several R packages in addition to CATALYST) or a file-size-limited online version. Links, installation instructions, example datasets and vignettes are accessible from the CATALYST project page: https://catalyst-project.github.io/.

## AUTHOR CONTRIBUTIONS

S.C., V.Z., M.D.R. and B.B. conceived the study. S.C. performed all single-cell experiments with help from V.Z. and H.C. S.E. performed the imaging experiments with help from V.Z. H.C. developed the CATALYST R package and the shiny app with the input from V.Z., M.D.R., S.C, and B.B. H.C, V.Z, S.C, and M.D.R. performed data analysis and interpretation. S.C. and H.C. wrote the manuscript with input from all authors.

## ACKNOWLEDGEMENTS

The authors thank Dr. Hartland Jackson for providing the breast cancer tissue sample (ethic approval ref. no. StV 12-2005), the Bodenmiller and the Robinson laboratories for fruitful discussions, and the CyTOF facility of the University of Zurich for providing us access to its instrument This work was supported by the Swiss National Science Foundation (SNSF) R’Equip grant, a SNSF Assistant Professorship grant PP00P3-144874, the PhosphonetPPM and MetastasiX SystemsX grant, and funding from the European Research Council (ERC) under the European Union’s Seventh Framework Programme (FP/2007-2013)/ERC Grant Agreement n. 336921. S.C. was funded by a Roche Postdoctoral Fellowship.

---

**Figure S1.**
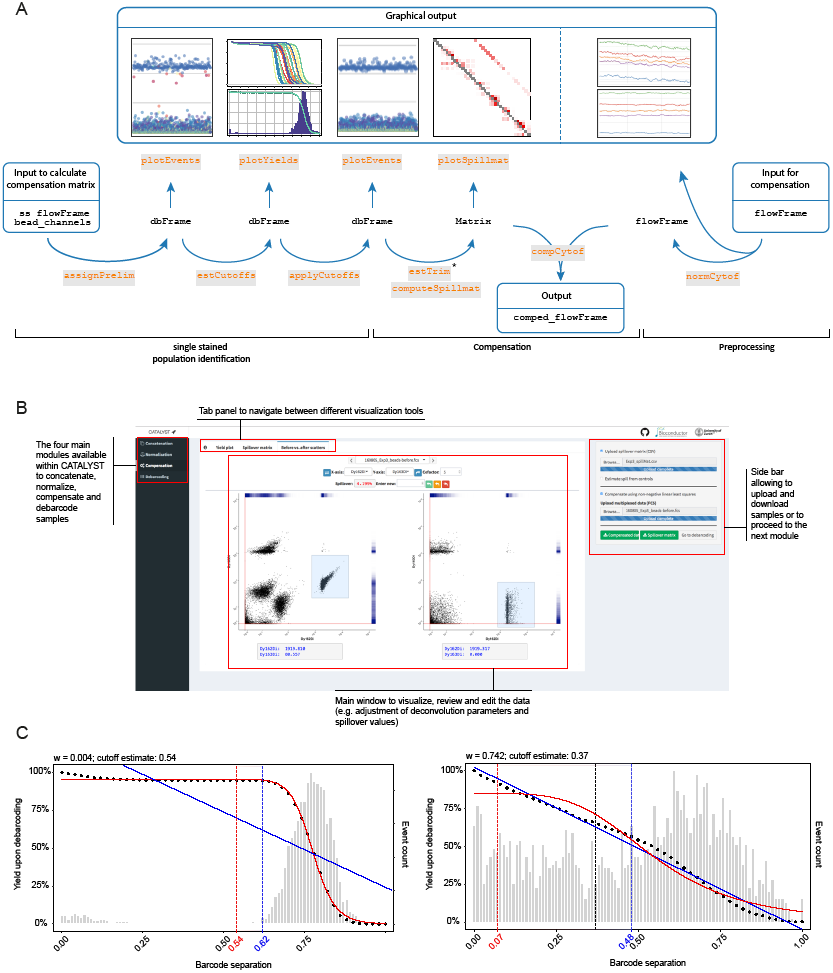
Description of the main functions of the CATALYST package. **A.** Schematic of the workflow used in the CATALYST package to generate a compensated file based on beads stained with single antibodies. The graphical outputs generated during the process are indicated above the steps. **B**. Screen shot depicting the main features available with the Shiny app. The compensation module is used as an example. **C.** Description of the automatic cutoff estimation for each individual population. The bar graphs indicate the distribution of cells relative to the barcode distance, and the dotted line corresponds to the yield upon debarcoding as a function of the applied separation cutoff. Data were fitted with a linear regression (blue line) and a three parameter log-logistic function (red line). The cutoff estimate is defined as the mean of estimates derived from both fits, weighted with the goodness of the respective fit (see Materials and Methods).

**Figure S2.**
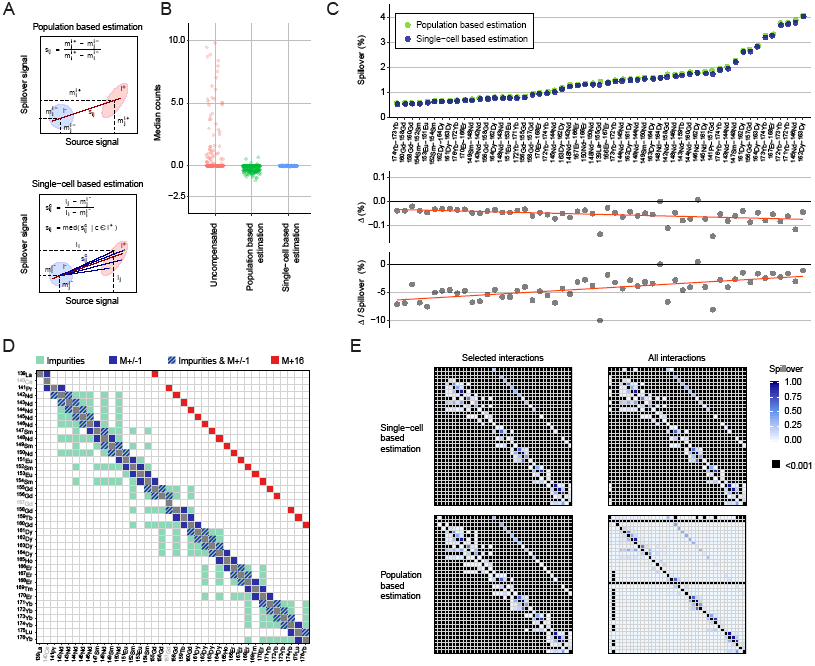
Description of the specificities of spillover matrix calculation for mass cytometry data. **A.** Scheme describing spillover estimates at the population level (upper panel) and at the single-cell level (lower panel). **B**. Dot plots showing the median counts in each channel potentially affected by spillover for beads compensated based on population estimates versus single-cell estimates compared to uncompensated data. **C.** Plots showing the spillover in percent for the main interactions as assessed at the population level and at the single-cell level (top panel) and the absolute difference (middle panel) and the relative difference (lower panel) in spillover percentages. **D.** Spillover matrix showing the interactions estimated by default in CATALYST. Only those interactions expected to occur based on impurities, abundance sensitivity, and oxidation are taken into consideration. **E.** Spillover matrix calculated for expected interactions versus all interactions using single-cell estimates (upper panels) versus population estimates (lower panels).

**Figure S3.**
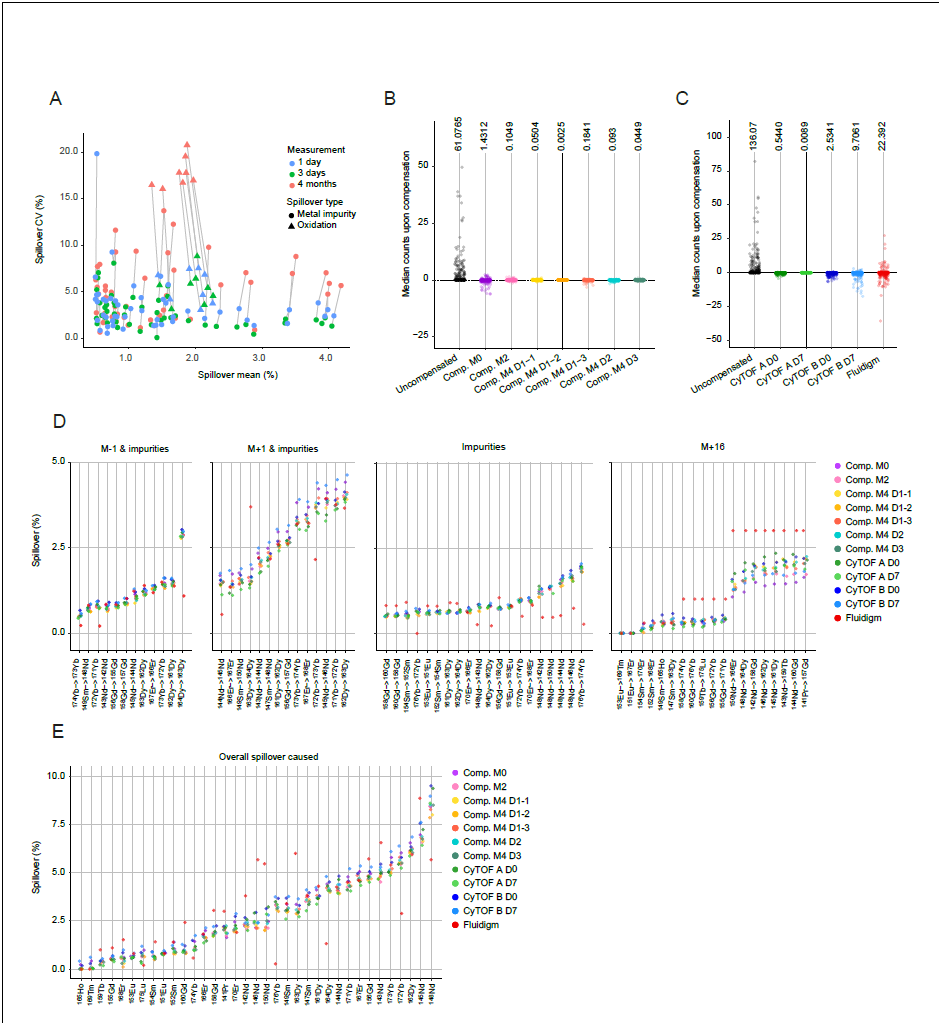
Compensation matrix stability over stainings, measurements, and instruments. **A.** Scatter plot displaying the means and standard deviations of the spillover measured the same day (blue), after three days (green), and after four months (red). The origin of spillover (metal impurity vs. oxidation) is indicated. **B.** Spillovers observed in single-stained beads in absence of compensation and upon compensation with each of seven different matrices acquired at the indicated time points are displayed as a dot plot. For each dataset, the average sum of squares is shown on top of the graph. **C.** Spillovers observed in single-stained beads without and upon compensation with each of four matrices acquired at the indicated time points on the indicated instruments are displayed as a dot plot. The compensation performed with the compensation matrix provided by Fluidigm is also shown. For each dataset, the average sum of squares is shown on top of the graph. **D**. Spillovers assessed for the individual relationships over stainings, measurements, and instruments are shown. Data are shown for M-1 and impurities (mean interaction > 0.5%), M+1 and impurities (>0.5%), impurities (>0.5%), and M+16 (all interactions). **E.** Scatter plot showing the total amount of spillover in each individual channel for the seven matrices acquired 4 months apart, for the four matrices acquired on two different machines, and for the theoretical matrix provided by Fluidigm.

**Figure S4.**
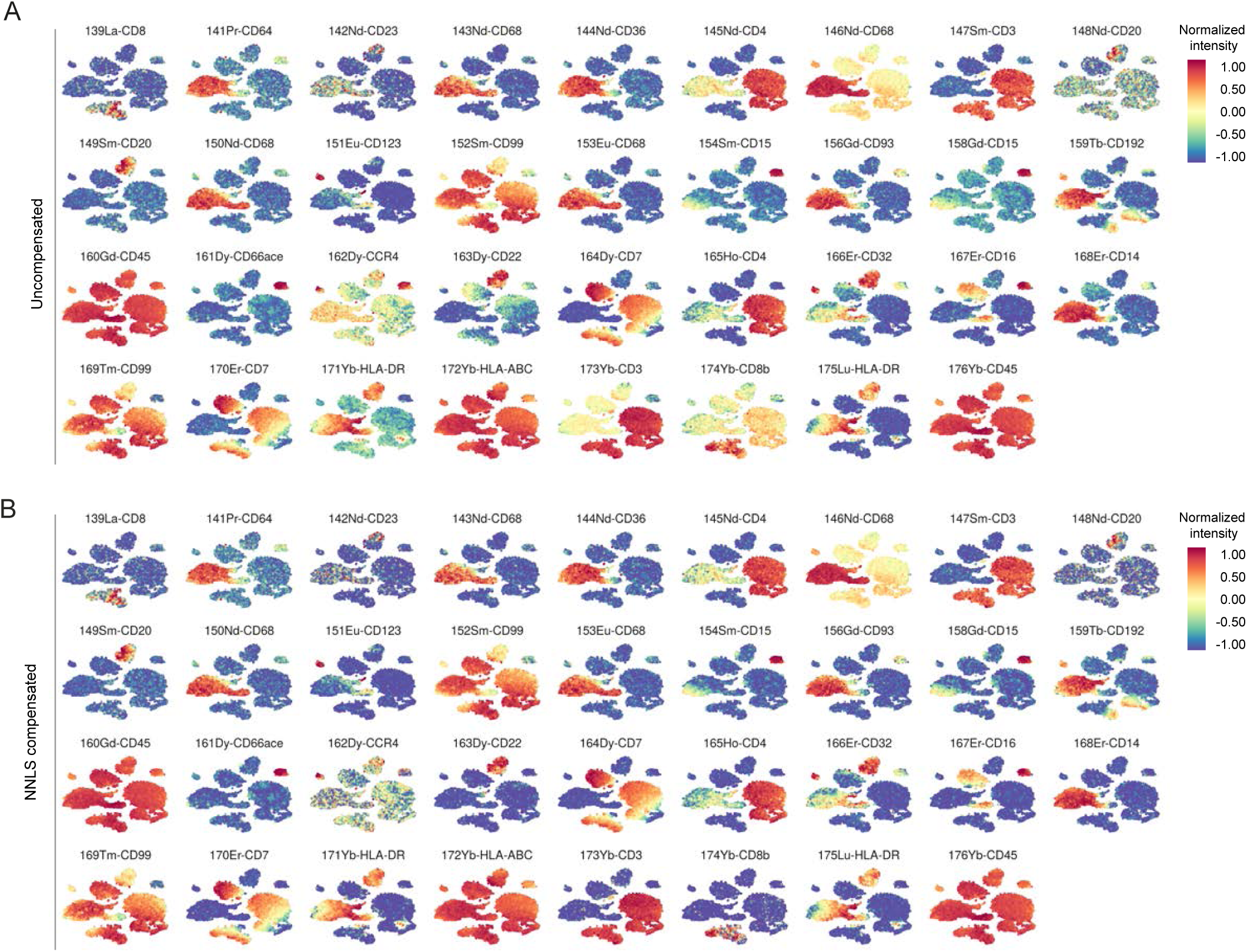
Compensation is necessary to ensure accurate interpretation of t-SNE maps. t-SNE maps displaying data on a subset of 20,000 PBMCs analyzed with our 36-antibody panel and colored by marker expression for all the antibodies included in the analysis (**A**) in absence of compensation and (**B**) after compensation based on NNLS.

**Figure S5.**
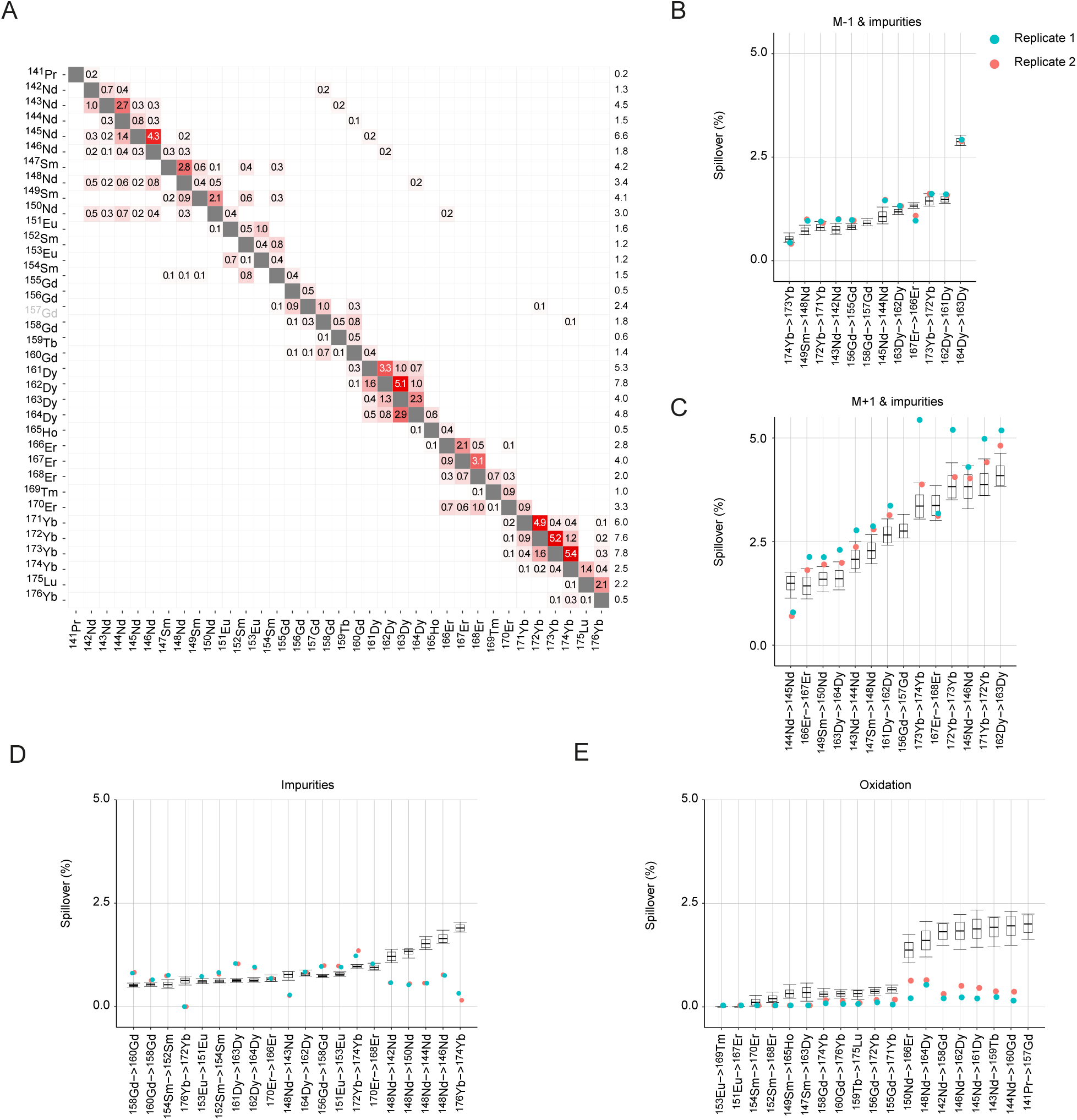
Compensation matrix for IMC. **A.** Spillover matrix calculated based on single isotope containing pixels. Values on the diagonals are one. Spillover is calculated only in potentially affected channels (Figure S2D). Numbers in the squares indicate percentages of spillover by channels in rows into channels in columns. Numbers in the last column show the total amount of spillover received in the corresponding channels. **B-E.** Signal interference for the indicated interactions shown for two independent IMC measurements of single isotopes spotted on a slide. Box plots show the spillover values obtained across the 11 replicates performed in flow mass cytometry, as described in Figure 5D.

## MATERIALS AND METHODS

### Cell preparation

PBMCs were isolated using histopaque (Sigma Aldrich) density gradient centrifugation of buffy coats from healthy donors obtained from the Zurich Blood Transfusion Service (www.zhbsd.ch). Cells at the interphase were harvested, washed twice in PBS, immediately fixed in 1.6% paraformaldehyde (Electron Microscopy Sciences) for 10 min at room temperature, and stored at −80 °C.

### Antibodies and surface staining

Provider, clone, and metal tag of each antibody used for suspension mass cytometry analysis are listed in Figure 2A. Antibody conjugations were performed using the MaxPAR antibody conjugation kit (Fluidigm) according to manufacturer’s instruction. After labeling, the concentration of each antibody was assessed using a Nanodrop (Thermo Scientific) and adjusted to 200 μg/mL in Candor Antibody Stabilizer. To determine the optimal concentration for PBMC staining, each conjugated antibody was titrated between 0.25 and 4 μg/mL. All antibodies used in this study were managed using the cloud-based platform AirLab^29^.

### Cell barcoding

To assess the effect of total metal load on spillover, 0.3-0.8 x 10^6^ cells from each tumor sample were barcoded using a 60-well barcoding scheme consisting of unique combinations of four out of eight barcoding reagents as previously described^14^. Six palladium isotopes (10^2^Pd, ^104^Pd, ^105^Pd, ^106^Pd, ^108^Pd, and ^110^Pd, Fludigm) were conjugated to bromoacetamidobenzyl-EDTA (BABE) and two indium isotopes (^113^In and ^115^In, Fludigm) were conjugated to 1,4,7,10-tetraazacy-clododecane-1,4,7-tris-acetic acid 10-maleimide ethylacetamide (mDOTA) following standard procedures^230^. For each concentration (20, 40, 80, 160, and 320 mM), cells were stained in triplicate using three random barcodes. Cells were barcoded using the transient partial permeabilization protocol described by Behbehani and colleagues^31^. Upon barcoding, cells were pooled and stained with the antibody mix.

### Cell and bead staining

Before antibody staining, cells were incubated with FcR blocking reagent (Miltenyi Biotech) for 10 min at 4 °C. One million of PBMCs were stained with 100 **u**l of the antibody mix (Figure 2A) for 30 min at 4 °C. Cells were washed twice in cell staining medium (CSM, PBS with 0.5% bovine serum albumin and 0.02% sodium azide) and resuspended in 1 ml of nucleic acid Ir-Intercalator (Fluidigm) in 1.6% PFA/PBS for 1 h at room temperature. Cells were then washed twice in PBS and twice in water. Before acquisition, cells were diluted to 0.5 x 10^6^ cells/ml in water. For bead-based compensation, aliquots of BD^TM^ Compbead Ig **K** beads (BD Biosciences) were stained individually with each of the antibodies used in the panel according to manufacturer’s instruction. Briefly, for each channel assessed in the panel, one full drop of BD^TM^ Compbead was loaded in a well of a v-bottom 96 well plate and stained with 1 μg of the corresponding metal-labeled antibody. Beads were stained for 15 min at room temperature. After staining, beads were washed three times in CSM and then pooled in a single tube. Beads were then fixed in 1.6% PFA/PBS for 1 h at room temperature. After fixation, beads were washed twice in PBS and twice in water. Before acquisition, beads were resuspended in 500 μL of water. Bead and cell data were acquired on a Helios mass cytometer (Fluidigm) using instrument-based dual-count calibration, noise reduction, and randomization. Cells were selected based on event length between 10 and 75 pushes. When required, exported flow cytometry standard (FCS) files were uploaded into Cytobank, populations of interest were manually gated, and events of interest were exported as new FCS files.

### IMC

To assess signal interference in IMC, metal isotopes were diluted to a concentration of 0.05 mM in Trypan Blue and arrayed on an agarose-coated microscopy slide. For each individual metal spot, an area of 400 x 5 pixels was ablated at a frequency of 200 Hz using the Hyperion mass cytometry system (Fluidigm)

For acquisition of multiplexed stained cells in imaging, breast cancer tissue sections (Ethic approval: StV 12-2005) were stained with a combination of anti-carbonic anhydrase-^166^Er (polyclonal, R&D Systems) and anti-KI67-^168^Er (8D5, CST) as previously described^19^. Upon staining, a region was analyzed by IMC using the Hyperion system (Fluidigm).

### Data analysis

#### Single-cell deconvolution

In order to identify single-positive populations from beads acquired as a pool, we applied the single-cell deconvolution (SCD) algorithm described in Zunder et al.^14^. In brief, events were preliminarily assigned to the sample for which their signal was strongest. Subsequently, doublet events (i.e., events whose separation between the primary channels and second highest signal fell below a threshold value) were excluded. We optionally allow for i) population-specific separation thresholds and ii) automated estimation of these thresholds. For the estimation of cutoff parameters, we considered yields upon debarcoding as a function of the applied cutoffs. Commonly, this function will be characterized by an initial weak decline, where doublets are excluded, and subsequent rapid decline in yields to zero. In between, low numbers of counts with intermediate barcode separation give rise to a plateau. As shown in Figure S1C, to facilitate robust estimation of an optimal cutoff, we fit a linear and a three-parameter log-logistic function^32^ to the yields function:

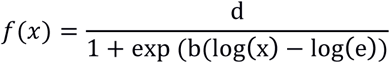

The goodness of the linear fit relative to the log-logistic fit is weighted as follows:

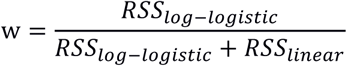

The cutoffs for both functions are defined as:

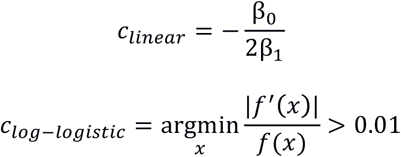

The final cutoff estimate *c* is defined as the weighted mean between these estimates:

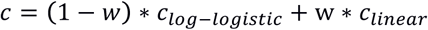

#### Estimation of the spillover matrix

To estimate the spillover matrix (*SM*), we made use of controls stained with individual antibodies. Because any signal not in a single-staining experiment’s primary channel *j* results from channel crosstalk, each spill entry *s*_*ij*_ can be approximated by the slope of a linear regression with channel *j* signal as the response, and channel *i* signals as the predictors, where *i* ∈ *w*_*j*_.

In a population-based fashion, this slope can be approximated as the ratio between the median signal of channel *i* positive events in channels *j* and *i*, 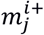 and 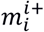, respectively. The expected background signal in these channels is computed as the median (or trimmed mean) signal of events that are i) negative in the channels *i* and *j* for which the spillover is investigated, ii) not assigned to interacting channels, and iii) not unassigned. These medians are indicated as 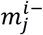 and 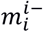, and subtracted, according to:

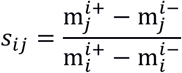

Due to mass cytometry data structure, we found that the following single-cell derived estimate is more accurate: Let *i*^+^ denote the set of cells that are positive in channel *i*, and 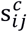 be the channel *i* to *j* spill computed for a cell c that has been assigned to this population. We approximate 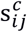 as the ratio between the signal in the unstained spillover receiving and stained spillover emitting channel, I_*j*_ and I_*i*_, respectively. Background signal is computed as above, and subtracted from all measurements:

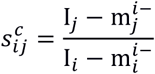

Each entry *s*_*ij*_ in *SM* is then computed as the median spillover across all cells c ∈ *i*^+^:

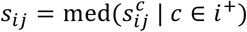

On the basis of their additive nature, spill values are estimated independently for every pair of interacting channels. By default, the current framework exclusively takes into account interactions that are sensible from a chemical and physical point of view: M±1 channels (abundance sensitivity), the M+16 channel (oxide formation), and channels measuring potentially contaminated metals (isotopic impurities). Optionally, all n·(n−1) possible interactions may be considered, and estimates below a threshold can be set to 0.

To generate the spillover matrix for imaging data, the single-stained images were imported into R and processed with the CATALYST package using individual pixels instead of individual cells as a readout.

#### Calculation of spillover and compensation

As demonstrated in Figure 1C, spillover is linear. In particular, the intensity observed in a given channel *j* is a linear combination of real signal and contributions from other channels that spill into it. If *s*_*ij*_ denotes the proportion of channel *j* signal that is due to channel *i* and w_j_ the set of channels that spill into channel *j*, then,

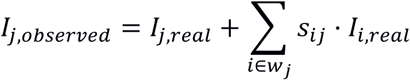

In matrix notation, measurement intensities may be viewed as the convolution of real intensities with a squared spillover matrix of dimensions p x p where p denotes the number of measurement parameters:

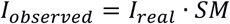

Note that diagonal entries *s*_*ii*_ = 1 for all *i* ∈ 1, …, *n*, where *n* denotes the number of measurement parameters, so that spill is relative to the total signal measured in a given channel. Assuming the correctness of this relationship, the resulting system of linear equations is traditionally solved exactly using linear algebra.

While mathematically exact, the solution to this equation does not account for measurement error or for the fact that the real signal would result in strictly non-negative counts. A simple and computationally efficient way to address this is to use non-negative least squares (NNLS)^25^. In brief, NNLS solves for *I*_*real*_ such that the least squares criterion is optimized under the constraint of non-negativity:

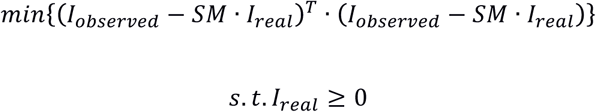

To arrive at such a solution we applied the Lawson-Hanson algorithm for NNLS as implemented in the ‘nnls’ R package.

For the image pixel compensation, the spillover matrix was exported as a tiff image and used for compensation using a custom written CellProfiler plugin (https://github.com/BodenmillerGroup/ImcPluginsCP)^28^. The images were visualized using ImageJ.

#### Segmentation and single-cell measurements

For segmentation, image stacks containing channels useful for segmentation were generated from the IMC raw data using the ‘imctools’ python package (https://github.com/BodenmillerGroup/imctools). The images were scaled up two fold using a CellProfiler pipeline and re-exported as tiff files suitable for Ilastik pixel classification. Using Ilastik the pixels of the image were classified as nuclei, cytoplasm/membrane, or background. The class information was exported as probability maps and used in CellProfiler for single-cell segmentation. The multiplexed images were measured in CellProfiler using a customized CellProfiler plugin (https://github.com/BodenmillerGroup/ImcPluginsCP). The single-cell data were then exported as csv files and imported into R for compensation with CATALYST and plotting.

### CATALYST R package

Installation instructions and open-source code are available through Bioconductor (http://bioconductor.org/packages/CATALYST). Detailed examples are included in the package vignette. Using flowCore^33^ infrastructure, CATALYST provides a user-friendly R implementation of the SCD algorithm. Furthermore, the package includes a function for estimation of the *SM* from *a priori* identified single-positive populations. The matrix returned by this workflow may be directly applied to the measurement data or exported for further use (e.g., to FlowJo or Cytobank).

